# *In vitro* Isolation and Identification of *Metarhizium pinghaense* Isolates and Assessment of Their Virulence against Whiteflies and Aphids

**DOI:** 10.1101/2025.11.19.689371

**Authors:** Haiyan Hu, Yali Wang, Chunyan Li, Ranran Zhang, Fangyu Liu, Xiaoan Sun

## Abstract

Four entomopathogenic fungal isolates were obtained from soil samples collected in Shouguang City, Shandong Province, using yellow mealworms as the baits. The result derived from the morphological and molecular identification of and assessment of the virulence against whiteflies (*Bemisia tabaci*) and the 4th instar aphids (*Aphis gossypii*) confirmed all four isolates as *Metarhizium pinghaense*, named SG-A, SG-B, SG-C, and SG-D, respectively. they were able to infect *B. tabaci* and *A. gossypii* with some differences in virulence and parasitic duration. The cumulative corrected mortality rates of *B. tabaci* and *A. gossypii* treated with the SG-A spore suspension (1×10^8^ conidia/ml.) were 94.44% and 96.67%, respectively in 8 days after inoculation. The LC_50_ and LT_50_ of SG-A against *B. tabaci* and *A. gossypii* were 7.00×10^4^, 4.21×10^5^ (conidia/ml), 4.13 and 2.61 (days) respectively. Moreover, SG-C was more pathogenic to *B. tabaci* than to *A. gossypii*, but SG-B and SG-D were less virulent to both insects. In conclusion, *M. pinghaense* SG-A is highly pathogenic and greatly lethal against both *A. gossypii* and *B. tabaci* and should be used as a potential biocontrol agent to control whiteflies and aphids at their nymph stage during the vegetable production season.

## Introduction

Whiteflies *Bemisia tabaci* (Gennadius) (Hemiptera: Aleyrodidae) and aphids *Aphis gossypii* (Glover) (Hemiptera: Aphididae) are important pests on over 1000 and 700 kinds of plant hosts and their host ranges are crossly overlapping, causing damages under suitable temperature and humidity year round ^[1–2]^. Moreover, as polyphagous and sucking pests, they feed on a wide range of plants, affect the vegetable growth and development and pose a serious economic threat through plant host saps and spreading more than 100 viruses such as watermelon mosaic virus (WMV), tomato yellow leaf curl virus (TYLCV), and tomato chlorosis virus (ToCV), devastating crop and vegetable production and yield^[3–6]^.

Currently, control of such insect pests by using chemical pesticides is effective in mitigating crop loss through pest prevention and management^[7–8]^. However, a long-term dependence on chemical control has carried a series of problems such as environmental pollution, plant resistance to insecticides (eg, pyrethroids, organophosphates, and neonicotinoids), and pesticide residues^[9–11]^. Therefore, it is urgent to find a green and efficient way of prevention and control. At present, biological control methods that utilize microbial parasitism or antagonism have gradually gained a great attention for their widely accepted applications in preventing and controlling of vegetable pests in facilities. Entomopathogenic fungi (EPF) *Metarhizium* spp. are important and potential substitutes for chemical pesticides to control pests. Of the 171 commercially available insect pathogenic fungi worldwide, approximately 34% of them belong to the genus *Metarhizium*^[12]^. The infection process of *Metarhizium* spp. is complicated, and can be divided into five stages: Recognition and attachment of conidia to the host insect skin, germination and formation of penetrating pegs to invade hemocoel, establishment and killing of the insect, and production of a large number of conidia^[13–14]^. *Metarhizium* spp. prove to be effective on a variety of insect pests, such as *Cacosceles newmannii*^[15]^, *Bagrada hilaris* (Hemiptera: Pentatomidae)^[16]^, and red palm weevil (*Rhynchophorus ferrugineus*) ^[17–18]^. However, environmental parameters, especially high temperature and humidity, greatly enhance the pathogenicity of those marketed *Metarhizium* spp. to infect and colonize their host insects^[19]^, which makes *Metarhizium* more feasible and effective in suppressing the population of both whiteflies and aphids in facility vegetable productions.

*Metarhizium* spp. are usually isolated from the rhizosphere in soil and utilized after initial *in vitro* screening and evaluation ^[20–21]^. The currently commercialized species of *Metarhizium anisopliae* has a wide range of host insects ^[20–23]^. At present, there are various isolates of *M. anisopliae* available in the market, but their biological control efficacy is unstable and their survival rate in field is unsatisfied. Moreover, there are few reports on the pathogenicity of *M. anisopliae* to sucking insect pests such as whiteflies and aphids and it is very much needed to isolate, screen, and evaluate new strains of wild-type *M. anisopliae*, which provides fundamental information and understanding for further research on their potential use in control of whiteflies and aphids in facility vegetable production in greenhouses. In this study, four *Metarhizium* isolates were obtained from soil samples in different regions, purified, identified as *M*. *pinghaense* by morphological and molecular biological methods, and evaluated for their pathogenicity against *B. tabaci* and *A. gossypii*. The results derived from this research should provide sufficient first-hand information on the biological control of sucking insect pests for the facility’s vegetable production.

## Materials and methods

### Fungal acquisition

Four fungal isolates were obtained from soil samples collected from vegetable-growing areas located at the Shouguang Modern Agriculture Center and Seed R&D Station with Weifang University of Science and Technology, Shouguang, Shandong Province (36°53’00.00’N, 118°42’00.00’E), China.

### Fungal isolation

Soil samples were collected 20 cm from below the surface and prepared for the isolation of Entomopathogenic fungal using yellow mealworm larva, *Tenebrio molitor* (Coleoptera: Tenebrionidae) as the baits^[24]^. Soil samples were poured onto a piece of clean newspaper, spread, and air-dried under laboratory conditions for 24 hours. 180 g of each soil sample was placed into a 250 mL sterilized flask with appropriate distilled water. Ten of the 6th-7th instar (1.2-1.8 cm long) healthy yellow mealworm larva were placed in each tissue culture bottle on the sample soil and kept in an artificial phenological incubator (spx) at (25 ± 2) ℃, RH 70% ± 5%, and photoperiod 12L:12D. All samples kept in the bottle were moisturize with an appropriate water spray and turned upside down at least 3 times a day without food supply throughout the incubation. The infected mealworms were removed from the soil and placed in a sterile 24-well plastic culture plate for further cultivation. The infected mealworm larvae were cut into small pieces, disinfected with 75% alcohol for 3-5 min, washed with sterile ddH_2_O for 5 times, and then placed onto SDAY nutrient plates (3 pieces of larvae per plate) containing 0.3 g/L tetracycline and 0.3 g/L kanamycin) for 10-14 days at 26 ℃. Fungal colonies were picked after 3–5 d growth and a single colony was obtained through transferring the edge of fungal colonies several times^[25–27]^.

### Identification of fungal

#### Morphological identification

A slide culture method was used by transferring a 5×5 mm PDA plug onto a sterilized slide and placing a small amount of purified fungal hyphae on the plug. The inoculated slide was covered and placed in a large Petri dish with a small amount of water in it, kept at 26℃ for 3 days and observed daily. The morphological characteristics of mycelia, conidiophores and conidia were observed under the microscope and photographed with the Motic Images Plus 2.0 system. The species were identified according to the morphological characteristics of the colony, mycelium, spore, and spore stalk^[25,28]^.

#### Molecular phylogenetic analysis of fungi

The total DNA of fungal isolates cultivated on PDA for 7 days was extracted by Norgen Biotek Fungi DNA Isolation Kit and the rDNA-ITS sequence of the strain was amplified by PCR using the universal primers ITS4/ITS1F. Also, Pbeta-F/Pbeta-R and PRPB2-F/PRPB2-R primers were used to amplify β-tubulin and RPB2, respectively (Table 1). The PCR reaction systems of DNA-ITS sequence, β-tubulin sequence and RPB2 sequence are: upstream and downstream primers (10mmoL) 1μL each, PCR Mastermix was12.5μL, template DNA1μL, ddH_2_O 9.5μL, a total of 25μL.

**Table 1.**
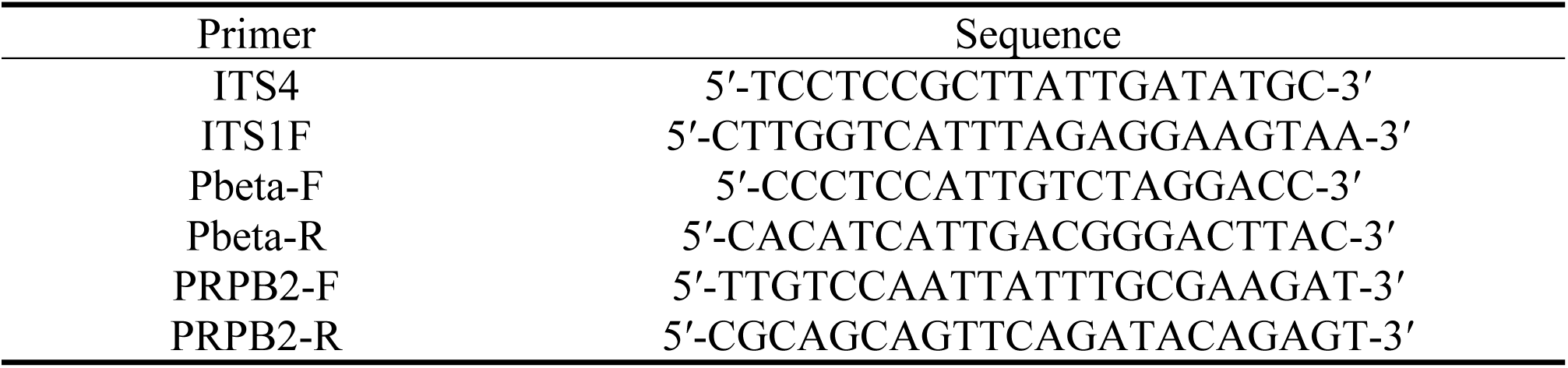
Primers used to identify the fungal isolates.

The PCR reaction procedures of the three sequences were: predenaturation at 95℃ for 3 min, denaturation at 94℃ for 1 min, annealing at 55℃ for 1min, extension at 72℃ for 1.5 min, 35 cycles. 1% agarose gel electrophoresis was performed and the amplification results were observed under the gel imaging system. PCR products were sent to Beijing Liuhe Bada Gene Technology Co., Ltd. for sequencing.

The sequences of rDNA-ITS, β-tubulin, and RPB2 were sequenced by the Blast program (http://blast.ncbi.nlm.nih.gov/Blast.cgi) and gene sequences in GenBank for homology analysis. The corresponding sequences of common related species were downloaded, and the corresponding sequences of *Beauveria bassiana* were taken as exogenous species. MEGA4.0 software ClustalX method was used to compare multiple sequences, and molecular phylogenetic trees were constructed by the neighbor-joining method. The system tree was tested with Bootstrap and repeated 1000 times.

### Insect rearing

*Bemisia tabaci* and *Aphis gossypii* adults were collected on pepper and cucumber plants, respectively from a greenhouse with the R&D station and raised on their host seedlings. The population of whitefly was identified as *Bemisia tabaci* Q biotype based on mitochondrial DNA COⅠ gene. After being reared for 5 generations, the 4th instar nymphs of both insects at the same age were used for the further experiments.

### Virulence bioassays

#### Preparation of conidial suspension

Four fungal isolates were grown on PDA plates for 14 days and the conidial spores were gently scraped off each of the colonies with soft-tipped sterilized spatula. Conidial spores were then rinsed once in a 200 ml beaker with 0.05% polyoxyethylene sorbitan monooleate solution (Tween 80, Sigma–Aldrich), resuspended in distilled water and mixed fully on a vortex shaker for 10 min. The average concentration of the spore suspension was determined by hemocytometer under the microscope for 5 times. Five concentrations (1×10^4^, 1×10^5^, 1×10^6^, 1×10^7^ and 1×10^8^ conidia/mL) of fungal spore suspension were prepared for further use. To determine the median lethal concentration of each fungal isolate to kill 50% of target insects hosts, (LC_50_), the suspension (1×10^8^ conidia/mL) was used to calculate the median killing time to kill 50% of the target insects (LT_50_).

### Insects bioassay

#### Laboratory bioassay of four fungi isolates against *B. tabaci* and *A. gossypii*

The bioassay was carried out following the method described by Paradza *et al.* (2021)^[29]^ with some modification. For each treatment, 30 4th-instar nymphs of *B. tabaci* or *A. gossypii* were placed onto a pepper leaf disc (3 cm in diameter) using a fine brush. The leaf discs with the nymphs were placed in a petri dish containing 10 ml 1% agar medium, sprayed separately with 2 ml different fungal spore suspensions in various concentrations (1×10^4^, 1×10^5^, 1×10^6^, 1×10^7^, and 1×10^8^ conidia/mL) using a small hand sprayer, air-dried, and reversely placed in an incubator (25±1℃, 70±5% RH and 12h light:12h dark photoperiod) for 8 days. Throughout the incubation, the infection status of the parasitized nymphs of whiteflies or aphids was observed daily and the hyphal growth of the parasitoid fungus and the fungal sporulation were recorded. A sterile water containing only Tween-80 was sprayed as a control, and the mortality rate of the control was kept below 10%. All treatments were replicated 3 times.

### Data analysis

The mortality rates were corrected using Abbott’s formula^[30]^. The Probit regression procedure was used to calculate the slope of the regress curve, intercept, 95% confidence limits, the lethal concentrations of 50% (LC_50_), and spore concentration of 1×10^8^ conidia/mL of median lethal time (LT_50_) of the isolates by SPSS 26.0 software, and calculate the regression equation based on the slope of the regress curve and intercept.

## Results

### Species identification of *Metarhizium* spp

#### Morphological identification

Four *Metarhizium* sp. isolates were collected from soil 20cm below the surface in different locations of the city. The pure culture of *Metarhizium* spp. was trapped by the yellow mealworms (Fig 1A-B) and transferred 3 times on cultivation plates. The fungal colony on the reverse side of SDAY Petri dish plates was reddish-brown, and folded longitudinally (Fig 2, A-1, B-1, C-1, D-1). Spores were yellow-green and oval, 2.01-2.58 × 5.84-6.25 µm (Fig 2, A-2, B-2, C-2, D-2). Under the microscope, hyphae were septate and the conidial peduncles were similar to the mycelium, with single or multiple small peduncles at the top, which were densely and neatly formed, and long chain-like conidia were formed at the end (Fig 2, A-3, B-3, C-3, D-3).

**Fig 1.**
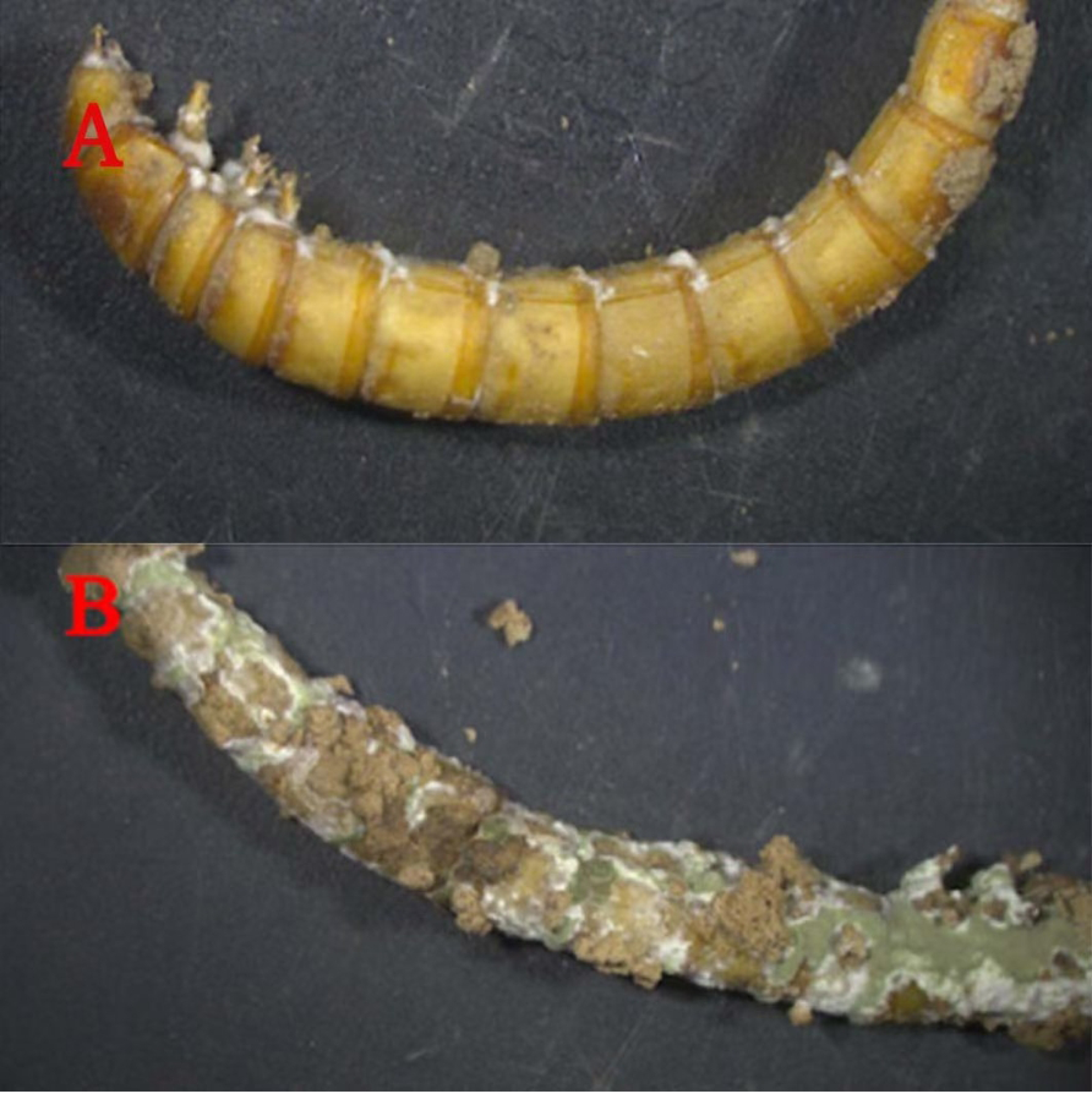
Appearance of mealworms infected and colonized by *Metarhizium sp.* in two (A) and four (B) days

**Fig 2.**
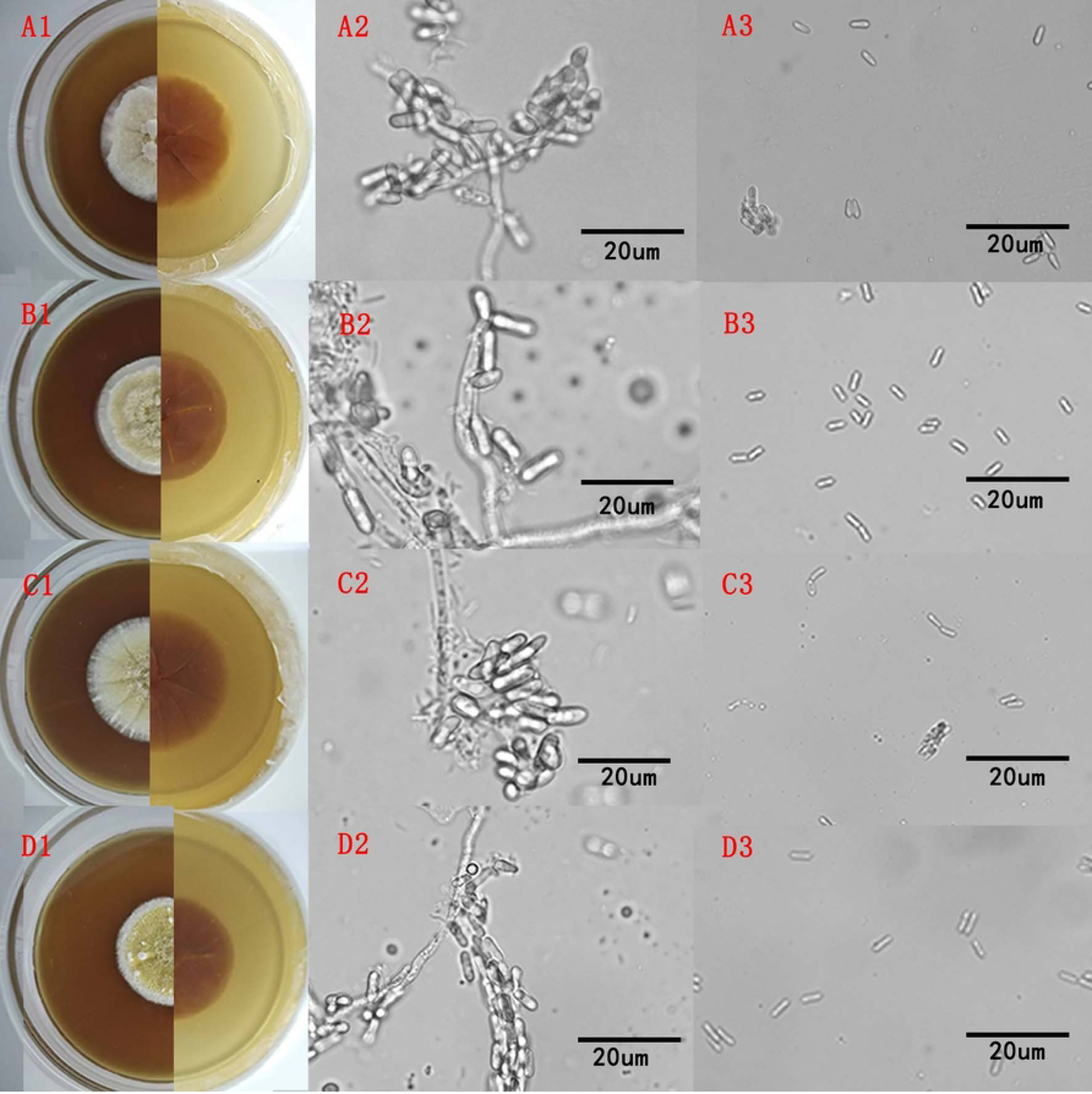
Morphological characteristics of *Metarhizium sp.* isolates (A-1, B-1, C-1, D-1), colonies of *Metarhizium sp.* cultured on SDAY at 25℃ for 14 days (A-2, B-2, C-2, D-2), and sporophore and conidial morphology of each isolate (A-3, B-3, C-3, D-3), respectively

#### Molecular biological identification

rDNA-ITS, β-tubulin, and RPB2 sequences were amplified with the DNA extracted from four *Metarhizium* isolates and the sequence alignment of 559bp, 1246bp, and 969bp fragments were respectively analyzed in GenBank by BLAST, showing a 99% genetic similarity with *Metarhizium pinghaense* with a constructed phylogenetic tree in comparison of the sequences from other *Metarhizium* species (Fig 3-5). Therefore, all of four isolates were identified as *M. pinghaense* according to their morphological traits and molecular identification.

**Fig 3.**
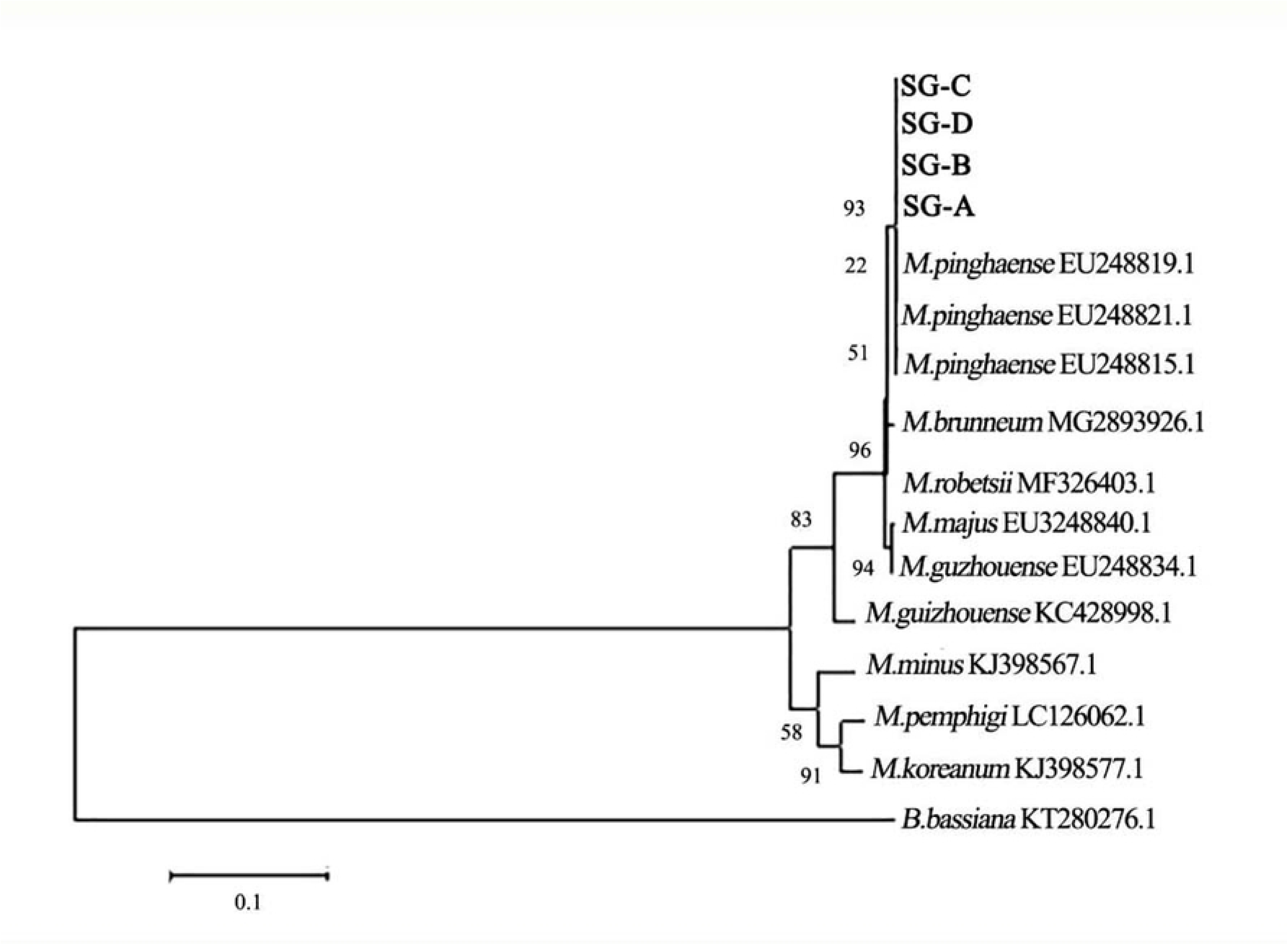
The phylogenetic tree of four isolates compared with other related *Metarhizium* species based on their sequence of the beta-tubulin coding gene through the neighbor-joining method

**Fig 4.**
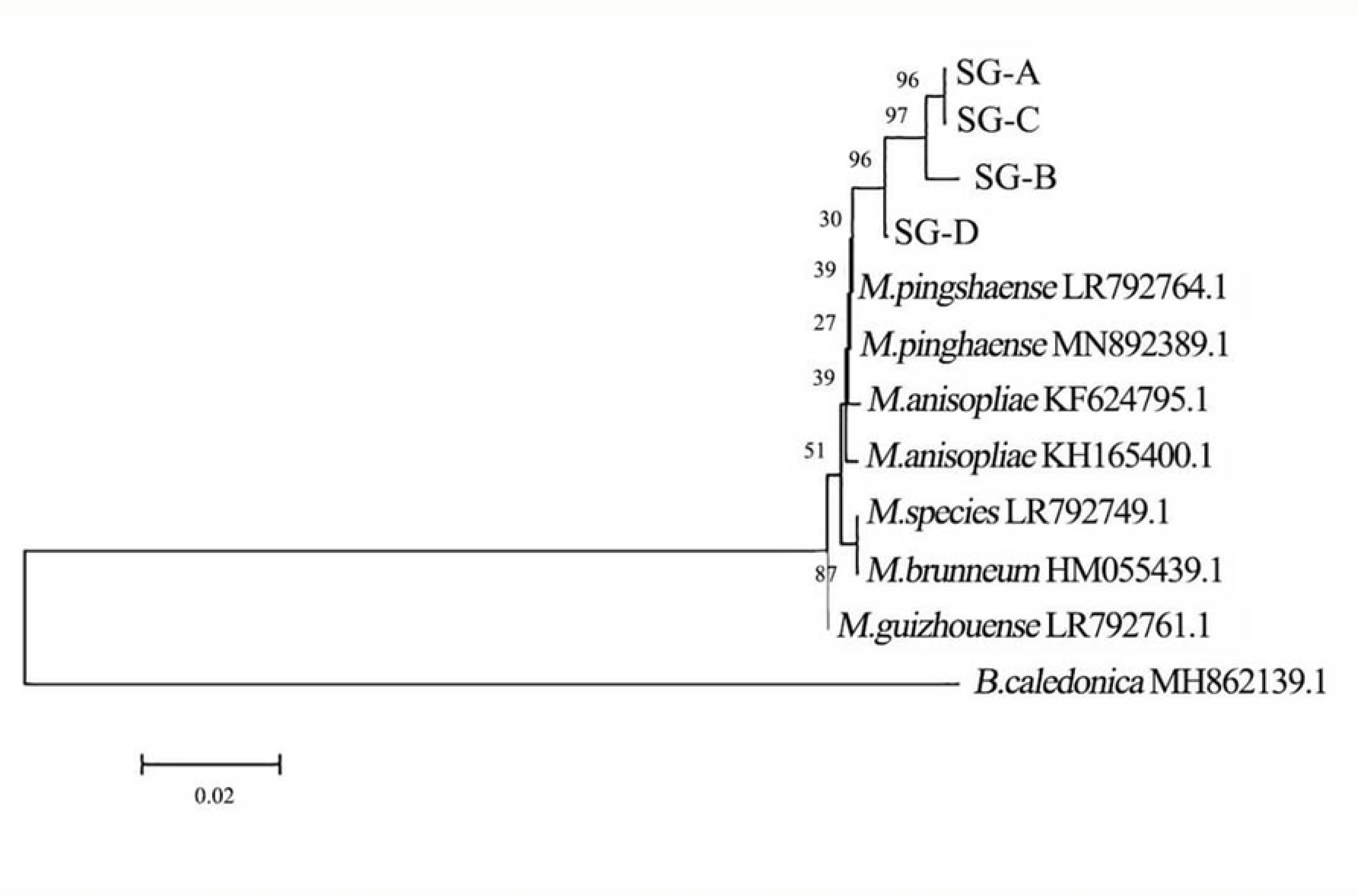
The phylogenetic tree of four isolates compared with other related *Metarhizium* species based on their sequence of the rDNA-ITS coding gene through the neighbor-joining method sequence

**Fig 5.**
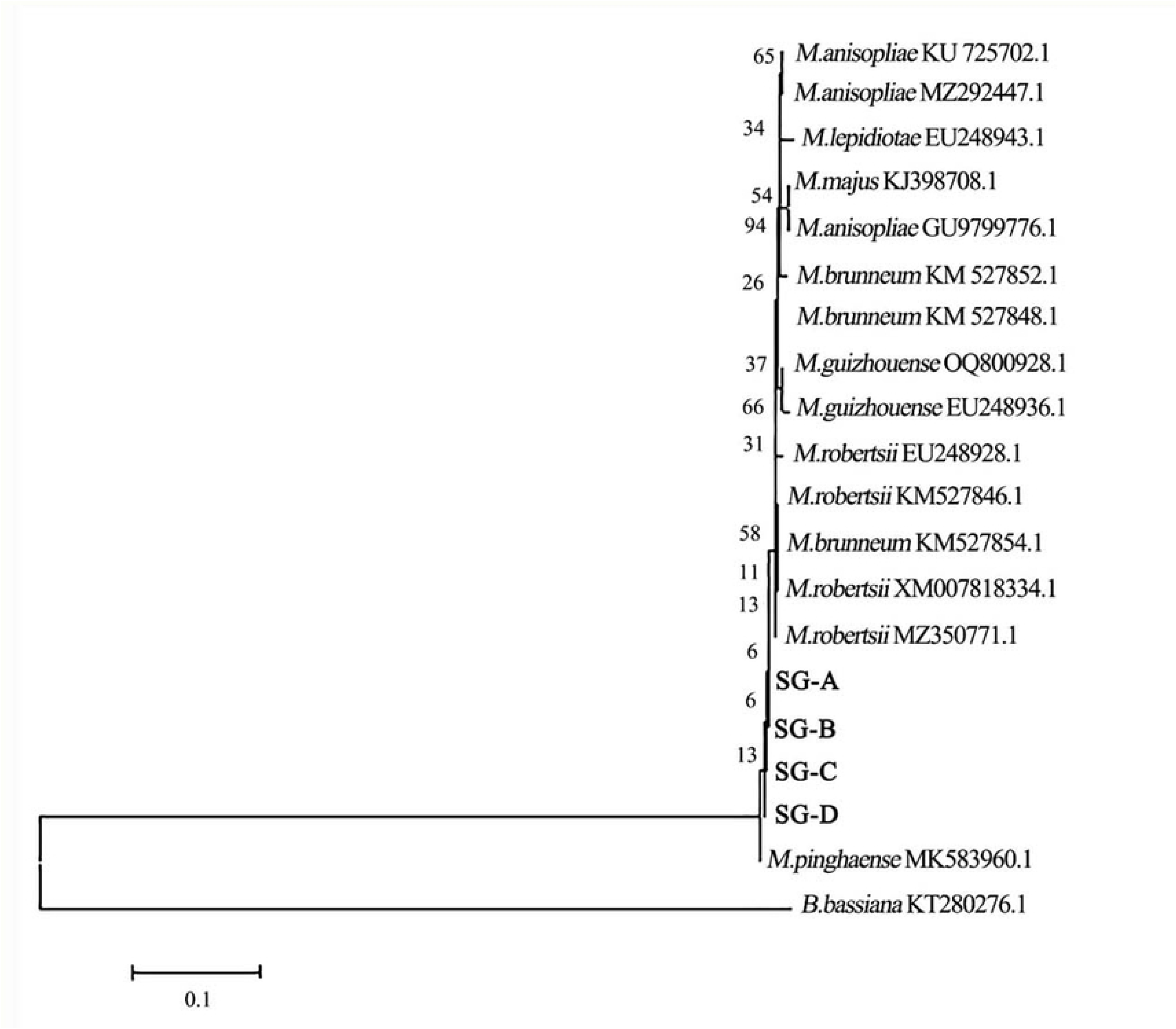
The phylogenetic tree of four isolates compared with other related *Metarhizium* species based on their sequence of the RPB2 coding gene through the neighbor-joining method

#### Bioassay

Two days after being submerged in the *M. pinghaense* spore suspension, the skin of the 4th instar whitefly nymphs turned dark with fluffy white fungal hypha (Fig 6 A). Four days after inoculation, the body of inoculated *M. pinghaense* nymph appeared to be yellow-green with heavy sporulating bodies (Fig 6 B).

**Fig 6.**
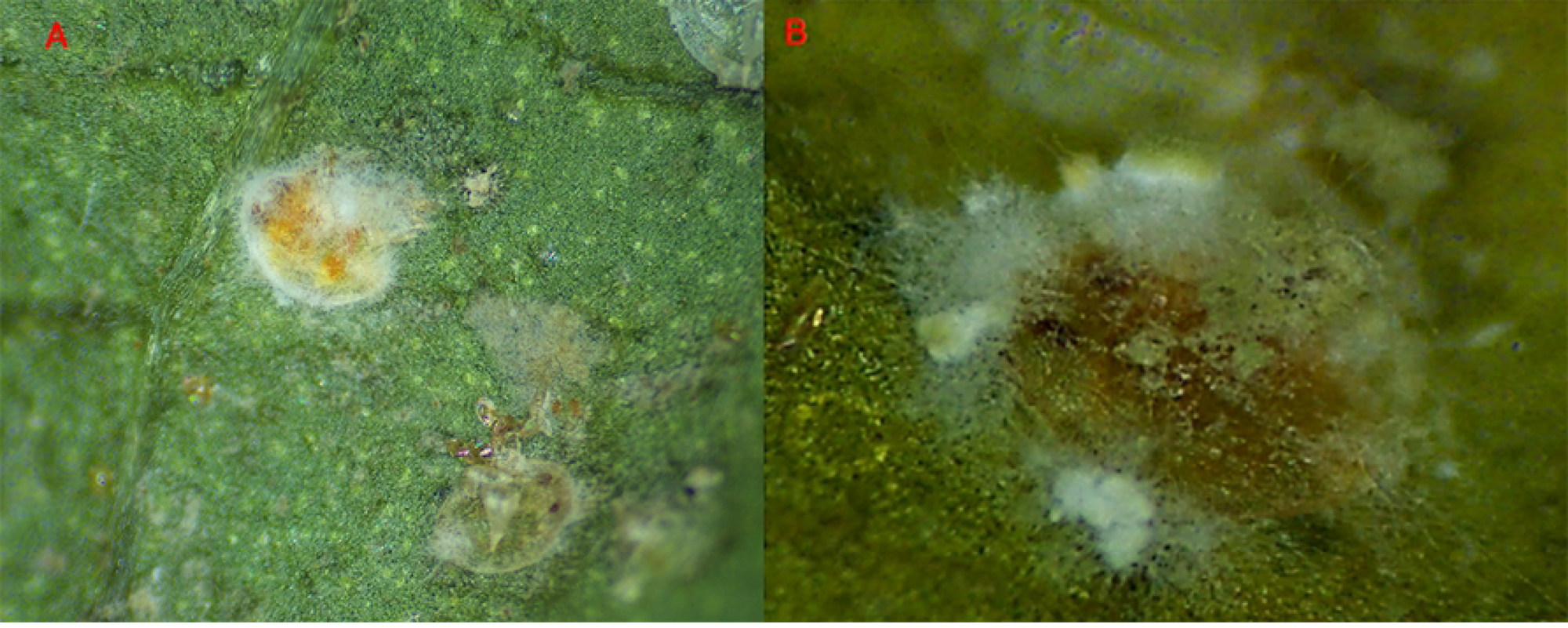
After treating the 4th instar nymphs of *B. tabaci* with a suspension of *M. pinghaense* spore, the dead nymphs were placed on a Petri dish lined with sterile moistened filter paper and incubated at 25 ± 1 ° C and >98% relative humidity for 2 days (A) and 4 days (B) to observe the symptoms of *M. pinghaens*e infection

Two days after being submerged in the *M. pinghaense* spore suspension, the skin of the 4th instar *A. gossypii* nymphs turned dark brown with light fungal mycelia (Fig 7A). Four days after inoculation, the body of the inoculated *M. pinghaense* nymph appeared to be black with heavy spore mass. The dead nymph bodies were covered with a massive chartreuse spores (Fig 7B).

**Fig 7.**
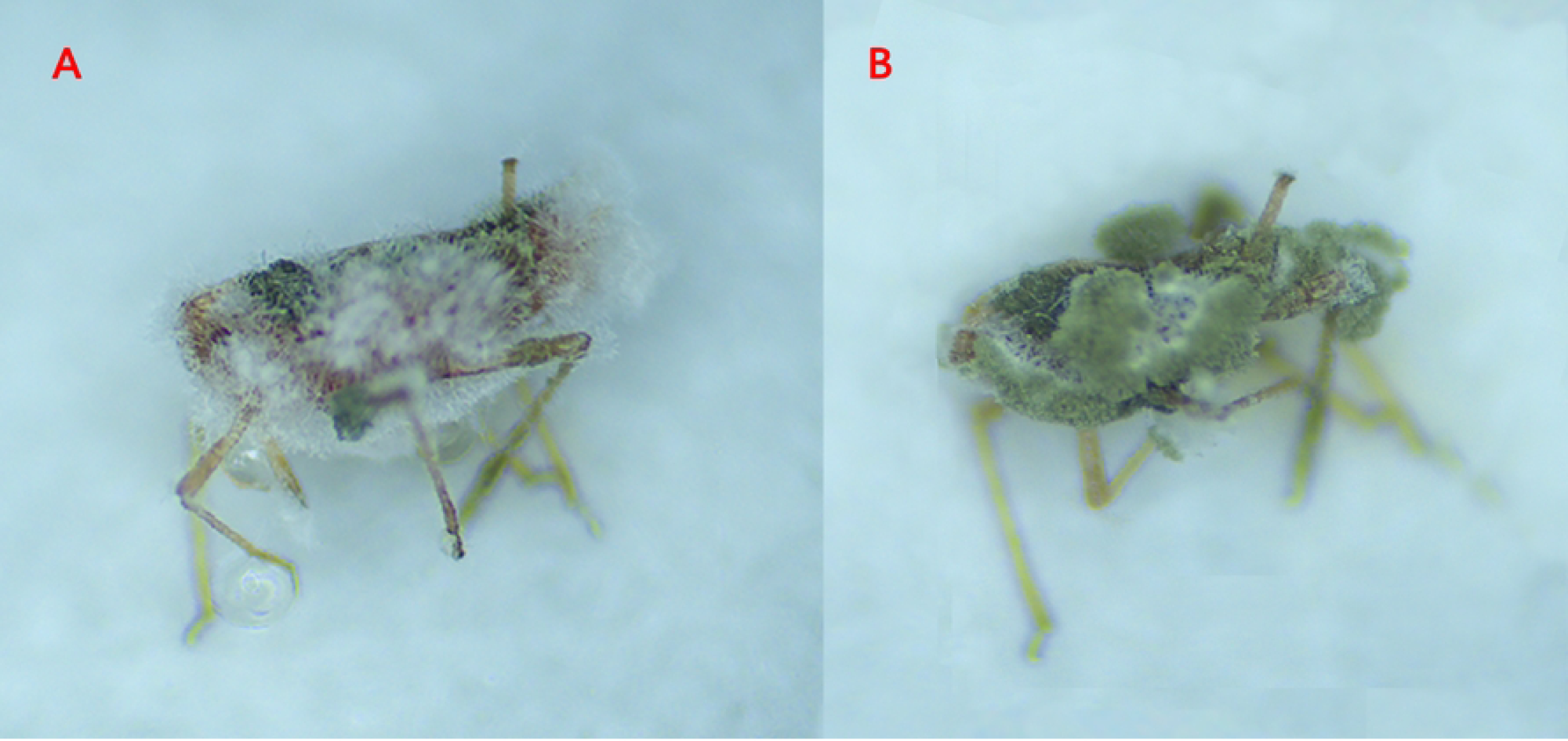
After treating the 4th instar nymphs of *A. gossypii* with a suspension of *M. pinghaense* spore, the dead nymphs were placed on a Petri dish lined with sterile moistened filter paper and incubated at 25±1℃ and >98% relative humidity for 2 days (A) and 4 days (B) to observe the symptoms of *M. pinghaense* infection

The 4th instar nymphs of whitefly inoculated 1×10^8^ conidia suspension of four *M. pinghaense* isolates started to die on Day 2 and the mortality increased exponentially through Day 5 (Fig 8). The *M. pinghaense* SG-A isolate scored the highest percentage causing whitefly nymphs to die (94.44% mortality in 8 days), followed by the SG-C, SG-B, and SG-D isolates (79.90% mortality in 8 days), respectively (Fig 8-A). Similarly, all 4 *M. pinghaense* isolates were lethal to *A. gossypii* nymphs 2 days after inoculation. The SG-A isolate showed the highest mortality (96.67% in 8 days), followed by SG-B, SD-D (Fig 8-B), and SG-C (58.89% in 8 days).

**Fig 8.**
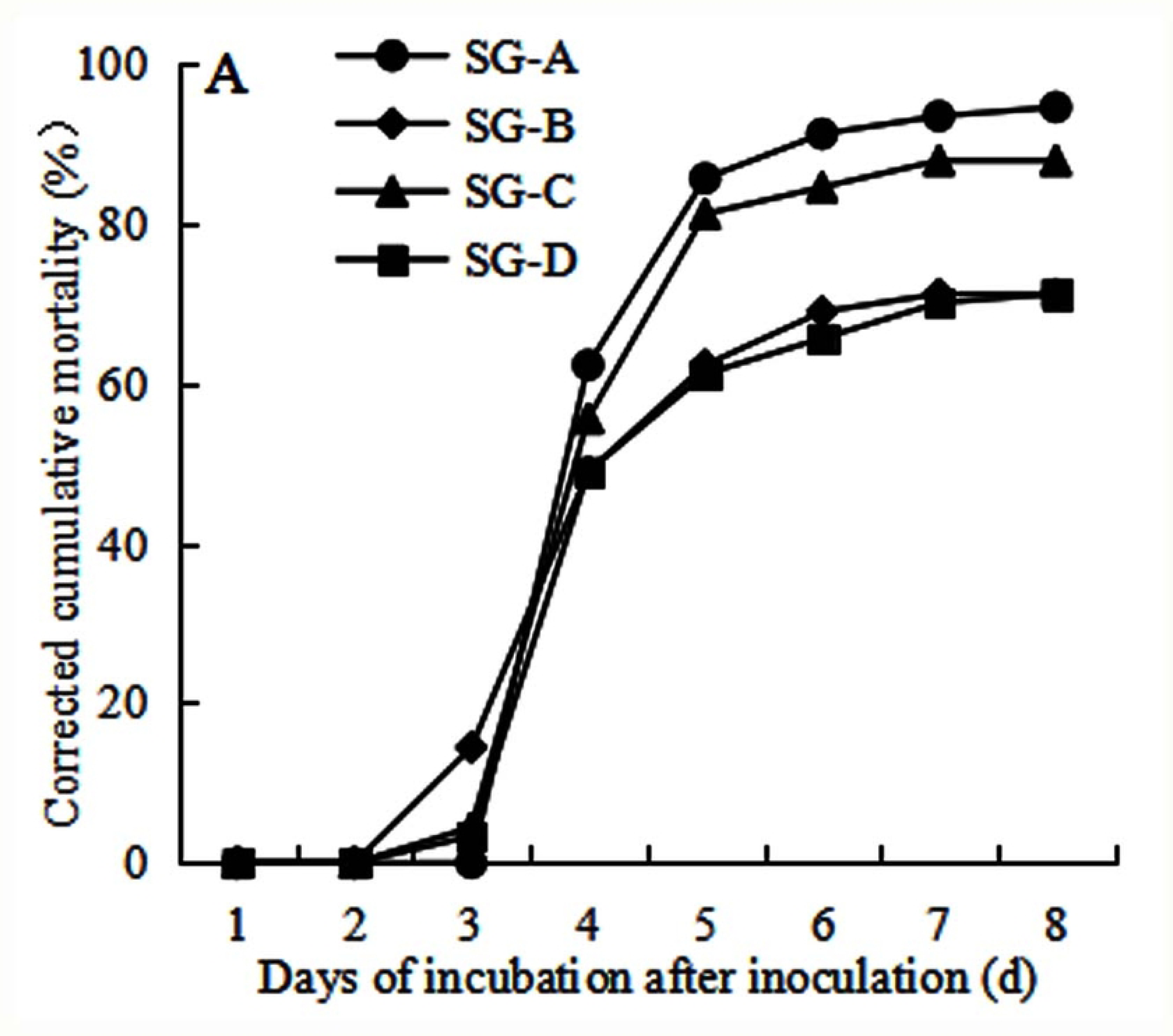
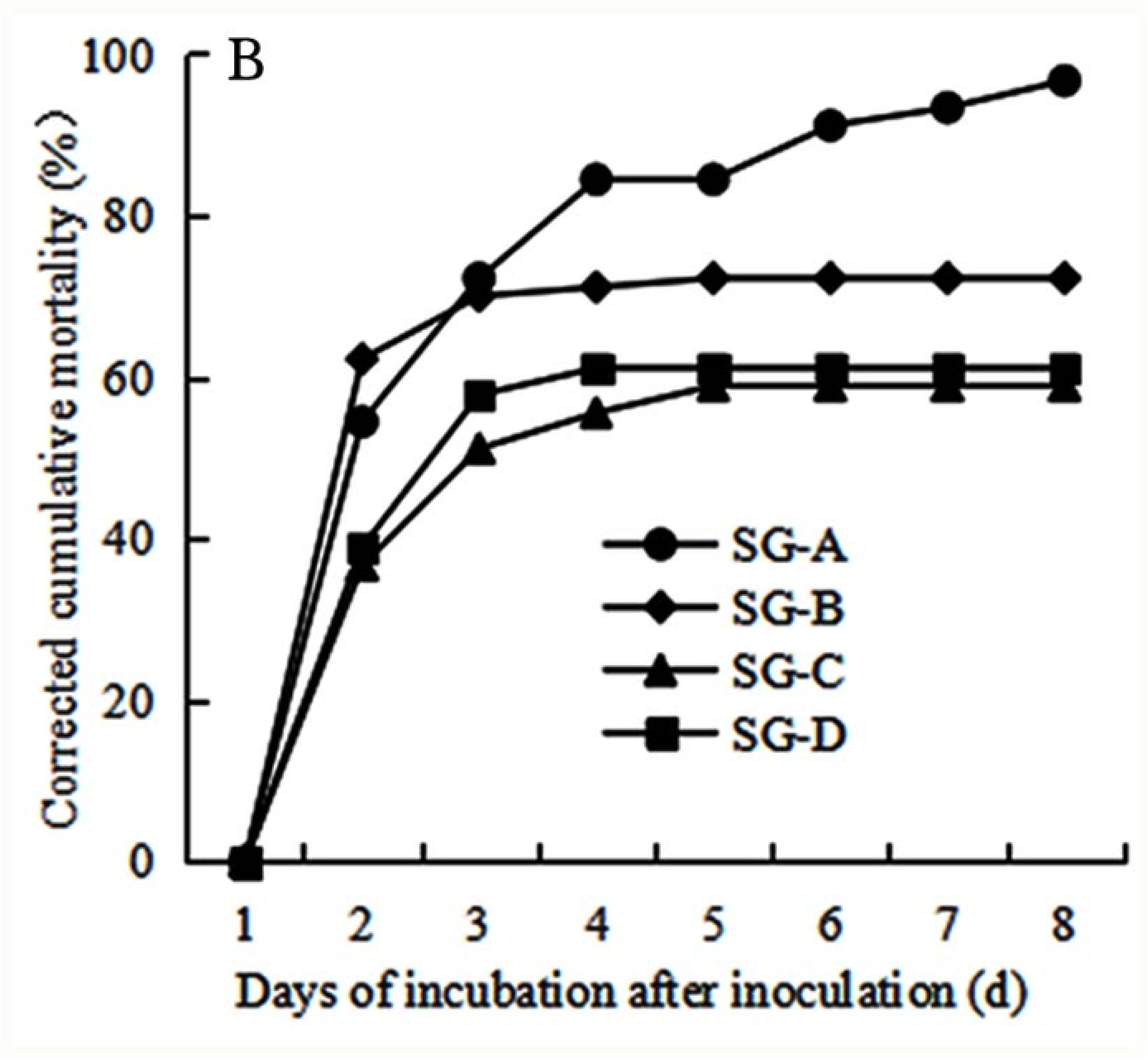
The mortality of the 4th instar nymphs of *B. tabaci* (A) and *A. gossypii* (B) over time after inoculation with four different *M. pinghaense* isolates at 1×10^8^ conidia/ml

There were differences in the mortality rates of *B. tabaci* and *A. gossypii* 8 days after infection by four strains of *M. pingshaense* at a spore concentration of 1×10 ⁸ conidia/ml. Among them, the SG-A strain exhibited the highest pathogenicity against both *B. tabaci* and *A. gossypii* (Table 2). After 8 days of treatment, the mortality rates of *B. tabaci* and *A. gossypii* nymphs were 94.44% and 93.33% respectively, showing significant differences from those of other strains (Table 2). In contrast, the pathogenicity of SG-C and SG-D strains against *B. tabaci* nymphs was higher than that against *A. gossypii* nymphs (Table 2). The SG-B strain showed comparable pathogenicity against both *B. tabaci* nymphs and *A. gossypii* nymphs (Table 2).There were differences in the median lethal time (LT_50_) of four *M. pinghaense* isolates against the 4^th^ instar *B. tabaci* nymphs treated with 1×10^8^ conidia/ml spore suspension. *M. pinghaense* SG-A only took 4.13 days on average to kill the wihtefly nymphs while the isolate SG-D needed 5.09 days on average to cause the death (Table 2). In terms of median lethal time (LT_50_) of four *M. pinghaense* isolates, SG-B took 2.31 days on average to kill its target nymphs, SG-C required at least 4.27 days on average to be lethal against *A. gossypii* nymphs at 1×10^8^ conidia/ml (Table 2).

**Table 2.** Cumulative mortality and median lethal time (LT50) after seven days post treatment of B. tabaci and A. gossypii nymphs after 8 days with spore suspension of four M. pinghaense isolates at 1×10^8^ conidia/ml.

| Isolate | % Mortality<br>of *B. tabaci*<br>(Mean±SE) | $LT_{50}$ (days)<br>(95%FL) | % Mortality<br>of *A. gossypii*<br>(Mean±SE) | $LT_{50}$ (days)<br>(95%FL) |
| --- | --- | --- | --- | --- |
| SG-A | 94.44±5.09a | 4.13(3.331-4.786) | 93.33±3.34a | 2.61(0.572-4.208) |
| SG-B | 71.11±6.94b | 4.87(4.377-5.415) | 72.22±5.09b | 2.31(1.927-2.659) |
| SG-C | 87.78±5.09a | 4.30(3.515-4.988) | 58.89±7.70c | 4.27(2.842-6.704) |
| SG-D | 71.11±5.09b | 5.09(4.168-6.201) | 61.11±3.85c | 3.85(2.238-6.185) |

The virulence of four *M. pinghaense* isolates against *B. tabaci* nymphs was quite different (Table 3). Among them, the isolate SG-A was more virulent (LC_50_ value of 7.00×10^4^ conidia/ml) than other isolates, followed by the isolate SG-C and SG-B. The least virulent SG-D had the LC_50_ value at 1.54×10^6^ conidia/ml. The virulence of four *M. pinghaense* isolates to the 4th instar nymphs of *A. gossypii* was also different (Table 3). The isolate SG-A caused the highest mortality on melon aphid nymphs (LC_50_ value at 4.21×10^5^ conidia/ml), followed by SG-B and SG-D. The lowest virulent isolate SG-C had a LC_50_ value at 1.58×10^7^ conidia/ml.

**Table 3.** The virulence of four *M. pinghaense* isolates against the 4th instar nymphs of *B. tabaci* and *A. gossypii*.

| Isolate | Insect species | Regression equation | LC<sub>50</sub> values ( conidia/ml ) | 95% confidence interval |
| --- | --- | --- | --- | --- |
| SG-A | *B. tabaci* | Y=-0.311 + 0.368X | 7.00×10<sup>4</sup> | 0.64×10<sup>4</sup>-2.81×10<sup>5</sup> |
|  | *A. gossypii* | Y=-0.646 + 0.398X | 4.21×10<sup>5</sup> | 9.38×10<sup>4</sup>-1.47×10<sup>6</sup> |
| SG-B | *B. tabaci* | Y=-0.451 + 0.211X | 1.36×10<sup>6</sup> | 8.17×10<sup>4</sup>-3.99×10<sup>7</sup> |
|  | *A. gossypii* | Y=-0.469 + 0.215X | 1.54×10<sup>6</sup> | 1.07×10<sup>5</sup>-4.61×10<sup>7</sup> |
| SG-C | *B. tabaci* | Y=-0.348 + 0.338X | 1.37×10<sup>5</sup> | 1.34×10<sup>4</sup>-5.80×10<sup>5</sup> |
|  | *A. gossypii* | Y=-0.978 + 0.306X | 1.58×10<sup>7</sup> | 3.06×10<sup>6</sup>-4.21×10<sup>8</sup> |
| SG-D | *B. tabaci* | Y=-0.469 + 0.215X | 1.54×10<sup>6</sup> | 1.07×10<sup>5</sup>-4.61×10<sup>7</sup> |
|  | *A. gossypii* | Y=-0.848 + 0.278X | 1.12×10<sup>7</sup> | 1.95×10<sup>6</sup>-3.91×10<sup>8</sup> |

## Discussion

The long-term and extensive use of chemical insecticides has raised serious issues such as pest resistance, food safety, environmental pollution, and health^[1,10]^. Therefore, it is imperative to apply the Integrated Pest Management (IPM) concept to minimize the amount and frequency of chemical pesticides and improve their efficacy as a safe and effective strategy in plant pest control. Biological control, especially through using parasitic fungi to control insect pests has been widely used. *Metarhizium* sp. is one of the most studied and wildly used biological control agents that can be easily isolated from the rhizosphere and has a potential to be used in the field after initial screening and evaluation with the advantages of being safe and adaptive in the facility environment^[31]^. While searching for the ideal candidate of *Metarhizium* as the biocontrol agent, the isolate identification is the first step and we used both morphological and molecular features to identify four *Metarhizium* isolates in this study. The morphological characteristics such as colony appearance, spore size, and spore shape of four *Metarhizium* isolates on SDAY plats were consistent with each other and to those described in previous studies^[22,32]^. To further ensure the accuracy of identification, molecular diagnosis is an additional and reliable method to determine the species, especially the alignment of key gene sequences after sequencing^[31]^. To date, partial multi-locus phylogenetic methods including ITS, TEF, RPB1, RPB2, and Pbeta gene sequences have gained popularity and recognition in describing the species within the *M. anisopliae* complex that includes nine confirmed species, including *M. pingshaense*^[32–33]^. The partial sequences of the ITS, Pbeta, and PRPB2 genes of 4 *Metarhizium* isolates were compared by BLAST with other same sequences of *Metarhizium* spp. reported in GeneBank and the constructed phylogenetic demonstrates that four isolates are grouped in a small evolutionary branch with a highest homology with *M. pingshaense* reported earlier. Along with the morphological characters, all of four strains are all *M. pingshaense*.

Early studies on the pathogenicity of *M. pingshaense* to other insects suggest that it could have infected many Lepidopteran species, such as rice leaf folder, *Cnaphalocrocis medinalis*^[23]^, yellow peach moth, *Conogethes punctiferalis*^[31]^ and Coleopteran insects including *Cacosceles newmannii*^[15]^, red palm weevil, *Rhynchophorus ferrugineus* (Olivier)^[22]^ with an acceptable efficacy.

Previous studies on the pathogenicity of parasitic fungi as biocontrol agent show that different species of insect parasitic fungi have various capabilities of infecting their target pests, even if they belong to the same species^[17]^. The results derived from our laboratory bioassays indicate that the mortality rates of four *M. pingshaense* isolates against *B. tabaci* and *A. gossypii* range from 71.1-94.4% and 58.9-96.7% at a concentration of 1×10^8^ conidia/mL, respectively, suggesting various degrees of pathogenicity or virulence against their target whiteflies and aphids. In addition, the study also indicates that the SG-A isolate shows the highest virulence against the 4th instar nympha of *B. tabaci* and *A. gossypii*, with a median lethal concentration to kill 50% of infected insects (LC_50_) with a suspension of 7.00×10^4^ conidia/mL and 4.21×10^5^ conidia/mL, respectively. The SG-A LC_50_ dosages to kill *A. gossypii* is higher than that to kill *B. tabaci*, possibly because the individual size of *A. gossypii* is bigger than that of *B. tabaci* for SG-A conidia to land and cause infections. The LT_50_ dosage of different isolates against *B. tabaci* and *A. gossypii* are different at the same concentration (1×10^8^ conidia/mL) of. The average of LT_50_ of four *M. pingshaense* isolates against *A. gossypii* takes a shorter time than that of those against *B. tabaci*, possibly due to the fact that aphids obtain more fungal spores on their appendages and body cavities through the crawling and are more susceptible to the fungal infections^[34]^. The SG-A isolate takes less time to kill *B. tabaci* with an LT_50_ at 4.13 d, while the SG-B does that on *A. gossypii* with LT_50_ at 2.31 d. Similar studies have shown that the isolates of *M. anisopliae*, AAUMB-29, AAUMFB-77, and AAUDM-43 have high pathogenicity against tobacco whitefly nymphs with an LC_50_ at 2.7×10^4^, 5.3×10^4^, and 5.4×10^4^ conidia/ml, respectively^[35]^. Norhelina’s research also demonstrates that the isolate GJ4 of *M. anisopliae* has the highest pathogenicity against tobacco whiteflies, with an LD_50_ at 6.62×10^4^ conidia/ml^[36]^.

Based on our results, we strongly believe that the isolate SG-A of *M. pingshaense* has effective pathogenicity against whiteflies and aphids and can be used as an excellent candidate to control both sucking insect pests. The screening and assessment of four *M. pingshaense* isolates under laboratory conditions not only provide an important theoretical basis for the subsequent prevention and control of whiteflies and aphids for facility vegetable cultivations, but also shed some light for further research and development on establishing their fermentation protocol and application.

## Supporting information

S1 Fig Overall situation of *M. pinghaense* infestation of aphids

S2 Fig Overall situation of *M. pinghaense* infestation of whiteflies

## Acknowledgments

Haiyan Hu and Yali Wang contributed equally to this work. The authors wish to thank Weifang Institute of Science and technology seed company and Facilities Vegetable pest prevention and control characteristic laboratory for technical supports.

